# Genetic effect of type 2 Diabetes to the progression of Neurological Diseases

**DOI:** 10.1101/480400

**Authors:** Md Habibur Rahman, Silong Peng, Chen Chen, Pietro Lio’, Mohammad Ali Moni

## Abstract

Neurological Diseases (NDs) are progressive disorder often advances with age and comorbidities of Type 2 diabetes (T2D). Epidemiological, clinical and neuropathological evidence advocate that patients with T2D are at an increased risk of getting NDs. However, it is very little known how T2D affects the risk and severity of NDs.

To tackle these problems, we employed a transcriptional analysis of affected tissues using agnostic approaches to identify overlapping cellular functions. In this study, we examined gene expression microarray human datasets along with control and disease-affected individuals. Differentially expressed genes (DEG) were identified for both T2D and NDs that includes Alzheimer Disease (AD), Parkinson Disease (PD), Amyotrophic Lateral Sclerosis (ALS), Epilepsy Disease (ED), Huntington Disease (HD), Cerebral Palsy (CP) and Multiple Sclerosis Disease (MSD).

We have developed genetic association and diseasome network of T2D and NDs based on the neighborhood-based benchmarking and multilayer network topology approaches. Overlapping DEG sets go through protein-protein interaction for hub protein identification and gene enrichment using pathway analysis and gene ontology methods that enhance our understanding of the fundamental molecular procedure of NDs progression.

Gene expression analysis platforms have been extensively used to investigate altered pathways and to identify potential biomarkers and drug targets. Finally, we validated our identified biomarkers using the gold benchmark datasets which identified the corresponding relationship of T2D and NDs. Therapeutic targets aimed at attenuating identified altered pathway could ameliorate neurological dysfunction in a T2D patient.

## I. Introduction

Diabetes mellitus(DM) derived from Greek word *Diabetes* stands *to pass through* and *Mellitus* is a Latin word for *honey*[1], often simply referred to as diabetes was firstly reported in Egyptian manuscript about 3000 years ago[2] and clearly made a distinction between type1 and type2 DM in 1936[3]. Type 2 Diabetes**(T2D)** mellitus first described as a component of metabolic syndrome in 1988[4] is the most common form of DM characterized by hyperglycemia, insulin resistance, and relative insulin deficiency. The global rise in type 2 diabetes (T2D) incidence estimating 439 million people by the year 2030[5] is known to be a nonreversible clinical issue with the attendant chronic kidney disease, amputation, blindness, various cardiac and vascular disease issues such *as strokes, heart disease, retinopathy, and peripheral ischemia. Its high incidence rate raises* issues of its interaction with other diseases occurring (or at risk of occurring) in the same individuals,i.e,co-morbidities. Epidemiological, clinical and neuropathological evidence advocate that patients with type 2 diabetes are at an increased risk of getting Neurological Diseases(NDs)[6,7,8]. Neurological diseases(NDs) are characterized by selective dysfunction and loss of neurons associated with pathologically altered proteins that deposit in the human brain but also in peripheral organs[7,9]. NDs primarily attack the neurons of the central nervous system and progressively damage the function of them. Neurons are most vulnerable to injury and normally don’t reproduce or replace themselves [10]. If neurons become damaged or die they cannot be replaced by medical treatments. So that NDs are very dangerous and currently they don’t have any cure. We studied several NDs Include Alzheimer Disease (AD), Perkinson Diseases (PD), Amyotrophic Lateral Sclerosis (ALS), Epilepsy Diseases(ED), Huntington Diseases (HD), Cerebral Palsy (CP) and Multiple Sclerosis Diseases (MSD) to find the effects of T2D on them.

Alzheimer Disease (AD), the most common type of incurable dementia is characterized by progressive neuronal loss, cognitive deterioration, and behavioral changes. Accumulation of amyloid or senile plaques and formation of neurofibrillary tangles are thought to be the major cause of neuronal loss in the AD brain[11,12]. PD is featured by progressive death of dopaminergic neurons in the substantia nigra and characterized by clinical symptoms including progressive impairment of movement control, cognitive decline, depression, anxiety, and olfactory dysfunction. Insulin dysregulation and changes in insulin action are concerned with developing Parkinson’s disease [11,13]. LGD also cognizant as Amyotrophic lateral sclerosis (ALS), is one of the major neurological diseases alongside Alzheimer’s disease and Parkinson’s disease that progressively damages motor neurons and muscle atrophy controlling voluntary muscle movement. Muscle weakness or stiffness is the initial symptoms of ALS. ALS patients are hypermetabolic and have impaired glucose tolerance associated with T2D[14]. ED is a heterogeneous group of neurodegenerative disorder that affects neural cells in the brain which are recognized by recurrent seizures or unusual behavior, awareness and sensations and high levels of β-amyloid in the brain can cause epileptiform activity[15]. HD is an inherited disorder characterized by the selective loss of medium spiny neurons from the striatum, leading to neuropsychiatric changes and movement disorder and the molecular cause of HD is the expansion of a CAG trinucleotide repeat in the gene encoding huntingtin(HTT), resulting in a polyglutamine stretch in the N-terminus of the protein[16]. Cerebral Palsy (CP) is a well-known group of disorders of movement, muscle tone, or other features that reflect abnormal control over motor function by the central nervous system[17]. Multiple sclerosis disease(MSD) is a chronic inflammatory and demyelinating disease of the central nervous system, which causes neurological disability due to the selective autoimmune destruction of white matter, ultimately leading to axonal loss. Genetic and environmental factors are thought to contribute to the pathogenesis of this disease[18][19].

How these are influenced by T2D is poorly understood, and typically studied by classical endocrinology approaches that focus on the effects of T2D associated cell secretions and serum glucose and glycation product levels. Although discoveries are continuing to be made in this field, the main causes or risk factors of NDs remain poorly understood.

To address these issues, here, we have proposed a novel computation-based approach, seeking to identify gene expression pathways common to T2D and NDs, as gene expression is profoundly affected by the disease processes themselves as well as predisposing genetic and environmental factors. Our approach aims to find overlapping pathways of potential clinical utility, but may also identify important new pathways relevance to many diseases.

With global transcriptome analyses, we investigated in detail common gene expression profiles of T2D and AD, PD, ALS, ED, HD, CP, and MSD. To understand the genetic effects of T2D on NDs, we examined gene expression dysregulation, disease association network, dysregulated pathway, gene expression ontology, and protein-protein interaction along with hub protein identification. We also investigated the validation of our study by using the gold benchmark databases (dbGAP and OMIM). This network-based approach identified significant common pathways influencing these diseases.

## II. MATERIALS AND METHODS

### A. Overview of analytical approach

A systematic and quantitative approach to evaluate human disease comorbidities using different sources of gene expression microarray data is summarized as shown in Fig. 1. This approach employs gene expression analyses, disease gene associations network, signaling pathway mechanism, Gene Ontology (GO) data, and protein-protein interaction network to identify putative components of common pathways between T2D and Neurological diseases (NDs).

**Fig. 1:**
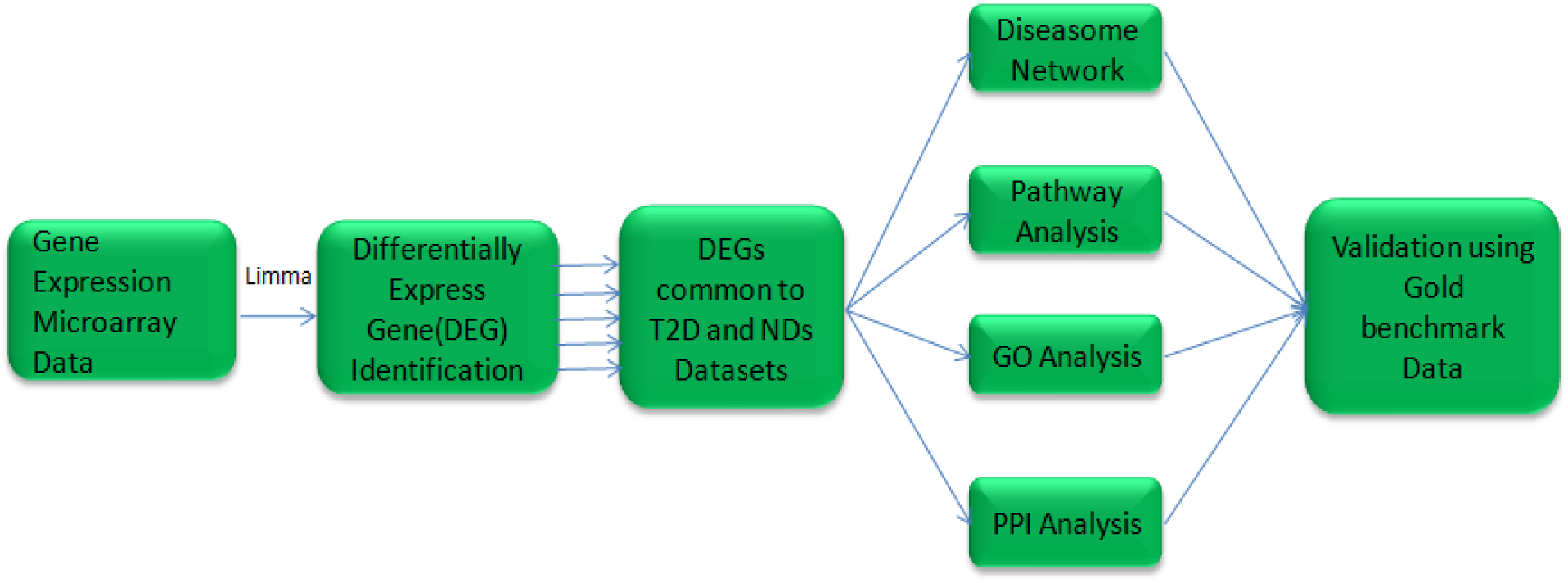
overview of the network-based approach.

### B. Datasets employed in this study

To investigate the molecular pathways involved in T2D on AD, PD, ALS, ED, HD, CP, and MSD at the molecular level, we first analyzed gene expression microarray datasets. In this study, we collected raw data from the Gene Expression Omnibus of the National Center for Biotechnology Information (NCBI) (http://www.ncbi.nlm.nih.gov/geo/).

We analyzed 8 different large human gene expression datasets having control and disease affected individuals for our study with accession numbers GSE23343[20], GSE28146[21], GSE19587[22], GSE833[23], GSE22779[24], GSE1751[25], GSE31243[26], and GSE38010[27]. The T2D dataset (GSE23343) contains gene expression data obtained through extensive analysis after conducting liver biopsies in the human. The AD dataset (GSE28146) is a microarray data on RNA from fresh frozen hippocampal tissue blocks that contain both white and gray matter, potentially obscuring region-specific changes. The PD dataset (GSE19587) is taken from 6 postmortem brains of PD patients and from 5 control brains. The LGD dataset (GSE833) is an Affymetrix Human Full Length HuGeneFL [Hu6800] Array. In this data, postmortem spinal cord grey matter from sporadic and familial ALGD patients are compared with controls. The ED dataset (GSE22779) is a gene expression profiles of 4 non-leukemic individuals (1 healthy and 3 with epilepsy) is generated from the mononuclear cells isolated from the peripheral blood samples before, and after 2, 6, and 24 hours of in-vivo glucocorticoid treatment. The CP (GSE31243) is an Affymetrix human genome U133A 2.0 array where 40 microarrays are provided into four groups to analyze the effect of cerebral palsy and differences between muscles. The Huntington Disease(HD) dataset(GSE1751) is an Affymetrix U133A expression levels for 12 symptomatic and 5 presymptomatic Huntington’s disease patients with 14 healthy controls. The MSD dataset (GSE38010) is a microarray data of multiple sclerosis (MS) patients brain lesions compared with control brain samples.

### C. Analysis methods

Analyzing oligonucleotide microarray data for gene expression is known to be an effective and responsive approach to demonstrate the molecular assessors of human diseases. Using DNA microarrays technologies with global transcriptome analyses, we compared the gene expression profiles of T2D with that in our AD, PD, ALS, ED, HD, CP, and MSD datasets to find the effect of T2D on them. All these datasets were generated by comparing diseased tissue with normal to identify differentially expressed genes (DEG) associated with their respective pathology. To uniform the mRNA expression data of different platforms and to avoid the problems of experimental systems, we normalized the gene expression data (disease state or control data) by using the *Z*-score transformation (*Z*_*ij*_) for each NDs gene expression profile using the following equation:

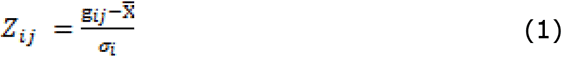

Where σ_*i*_ implies the standard deviation, *g*_*ij*_ represents the value of the gene expression *i* in sample *j.* These datasets were obtained and detected DEGs with their respective pathology to compare disease tissue with a normal one. We used *log*_*2*_ transformation and calculated expression level of each gene implementing linear regression: *Y*_*i*_ = a_o_ + a_1_*X*_*i*_ where *X*_*i*_ is a disease state(case and control),*a*_*0*_ and *a*_*1*_ *are* considered the coefficient of this model estimating with least squares and *Y*_*i*_ is the gene expression value. Student’s unpaired t-test was employed between two conditions and performed to find out DEGs over normal samples in patients and significant genes were preferred. A threshold of at least 1 log2 fold change and a p-value for the t-tests of <= 5 ∗ 10-2 were chosen. In addition, a two-way ANOVA with Bonferroni’s post hoc test was used to determine statistical significance between groups (<0.1). Gene symbol and title of different genes are extracted from each disease. Null gene symbol records are discarded from each disease. We also find out unique genes both over and under-expression genes. The most important up and down-regulated genes are selected between individual disease and T2D.

There are applied neighborhood benchmarking and topological methods to show the associations between genes and diseases. A gene-disease network was built(GDN) regarding the connection of gene-disease where nodes can be either diseases or genes such a network can be represented as a bipartite graph whether T2D is the center of this network using Cytoscape V 3.6.1[28]. Diseases are associated when they share at least one significant dysregulated gene. Given a particular set of human diseases *D* and a set of human genes *G*, gene-disease associations attempt to find whether gene *g* ∈ *G* is associated with diseases *d*∈ *D*. If *G*_*i*_ and *G*_*j*_, the sets of significant up and down-dysregulated genes associated with diseases *D*_*i*_ and *D*_*j*_ respectively, then the number of shared dysregulated genes 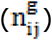 associated with both disease *D*_*i*_ and *D*_*j*_ is as follows[29]:

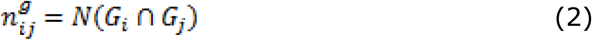

Co-occurrence refers to the number of shared genes in the GDN and common neighbors identified employing the Jaccard Coefficient method, where the edge prediction score for the node pair is:

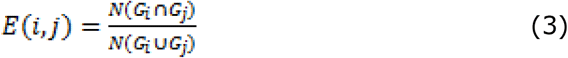

Where G is the set of nodes and E is the set of all edges. We used R software packages “comoR”[30] and “POGO”[31] to cross check the disease comorbidity associations.

To get further insight into the molecular pathways of T2D that overlap with the AD, PD, ALS, ED, HD, CP, and MSD, we performed pathway and gene ontology analysis using the Enrichr[32], and KEGG pathways database [33]. We also generated a protein-protein interaction (PPI) network for each disease pair datasets, using data from the STRING database [stringdb.org] where proteins are represented by nodes and the interaction between two proteins are represented by edges. Furthermore, we also incorporated three gold benchmark verified datasets, OMIM (www.omim.org), OMIM Expanded, and dbGaP (www.ncbi.nlm.nih.gov/gap) for retrieving the genes of all known diseases, relevant disorders, and genotype-phenotype relationships, in our study for validating the proof of principle of our network-based approach.

## III. RESULTS

### A. Gene Expression Analysis

To investigate the effects of T2D on NDs, we analyzed the gene expressing microarray data from the National Center for Biotechnology Information (NCBI) (http://www.ncbi.nlm.nih.gov/geo/). We found that 1320 genes were differentially expressed for T2D with False Discovery Rate(FDR) below 0.05 |logFC| >= 1 using R Bioconductor packages(Limma). Among them, 622 and 698 were up and down-regulated respectively. Similarly, our analysis identified the most significant differentially expressed genes for each NDGD after various steps of statistical analysis. We identified differentially expressed genes, 1606 (847 up and 759 down) in AD, 1588 (1167 up and 422 down) in PD, 2901 (735 up and 2166 down) in ALS, 1887 (882 up and 1007 down) in ED, 1338 (365 up and 973 down) in HD, 588 (243 up and 345 down) in CP and 7463 (3987 up and 3476 down) in MSD. The cross-comparative analysis was also performed to find the common differentially expressed genes between T2D and each ND. We observed that T2D shares 3,21,4,13,7,11 and 35 significantly up-regulated genes and 12,2,25,16,6,5 and 28 significant down-regulated genes with AD,PD,ALS,ED,HD,CP and MSD respectively. To find statistically significant associations among these diseases, we built up and down diseasome relationships network centered on the T2D as shown in fig. 2A and 2B in which two diseases are comorbid if there exist one or more genes that are associated with both diseases [34].Notably, 5 significant genes FLI1, PACSIN2, BICD1, TCP11L2, and ENTPD1 are commonly up-regulated among T2D, ED, and MSD. While 2 significant genes PEG10 and EFCAB14 are up-regulated among T2D, PD, and MSD. ITGB8 is up-regulated among T2D, HD, PD, and MSD.FBLN1 is up-regulated among T2D, ALS, and MSD.IGFBP5 is up-regulated among T2D, CP, and MSD. SGCB is up-regulated among T2D, HD, and MSD.SLC25A30 is up-regulated among T2D, PD, and CP. On the other hand, ST6GALNAC5, RIMS1 are down-regulated among T2D, AD, and MSD.ZBTB7A, YME1L1 are down-regulated among T2D, ALS, and ED. FUT6 is down-regulated among T2D, AD, and ALS.BRF1 is down-regulated among T2D, ALS, and CP.CDC14B is down-regulated among T2D, ALS, ED, and MSD.CD47 is down-regulated among T2D, ALS, HD, and ED.NRG1 is down-regulated among T2D, ALS, HD, and MSD.DNM1 is down-regulated among T2D, CP, and MSD.TLB1XR1 is down-regulated among T2D, HD, and MSD.GPR161 is down-regulated among T2D, ALS, and MSD.

**Fig. 2A.**
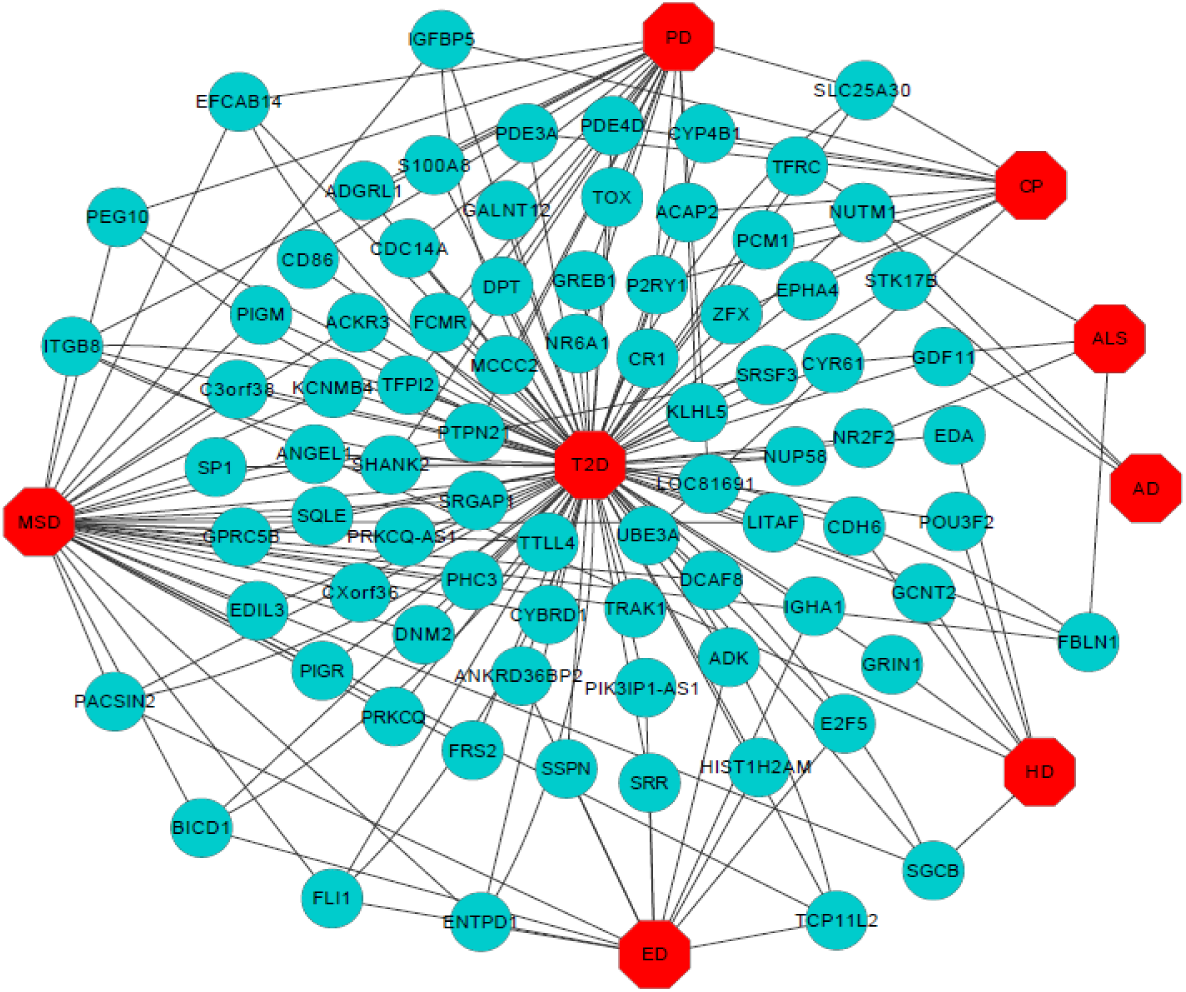
Disease network of T2D with NDs. Red colored octagon-shaped nodes represent different categories of disease, and round-shaped…… colored nodes represent commonly up-regulated genes for T2D with the other neurodegenerative disorders. A link is placed between a disease and a gene if mutations in that gene lead to the specific disorder.

**Fig. 2B.**
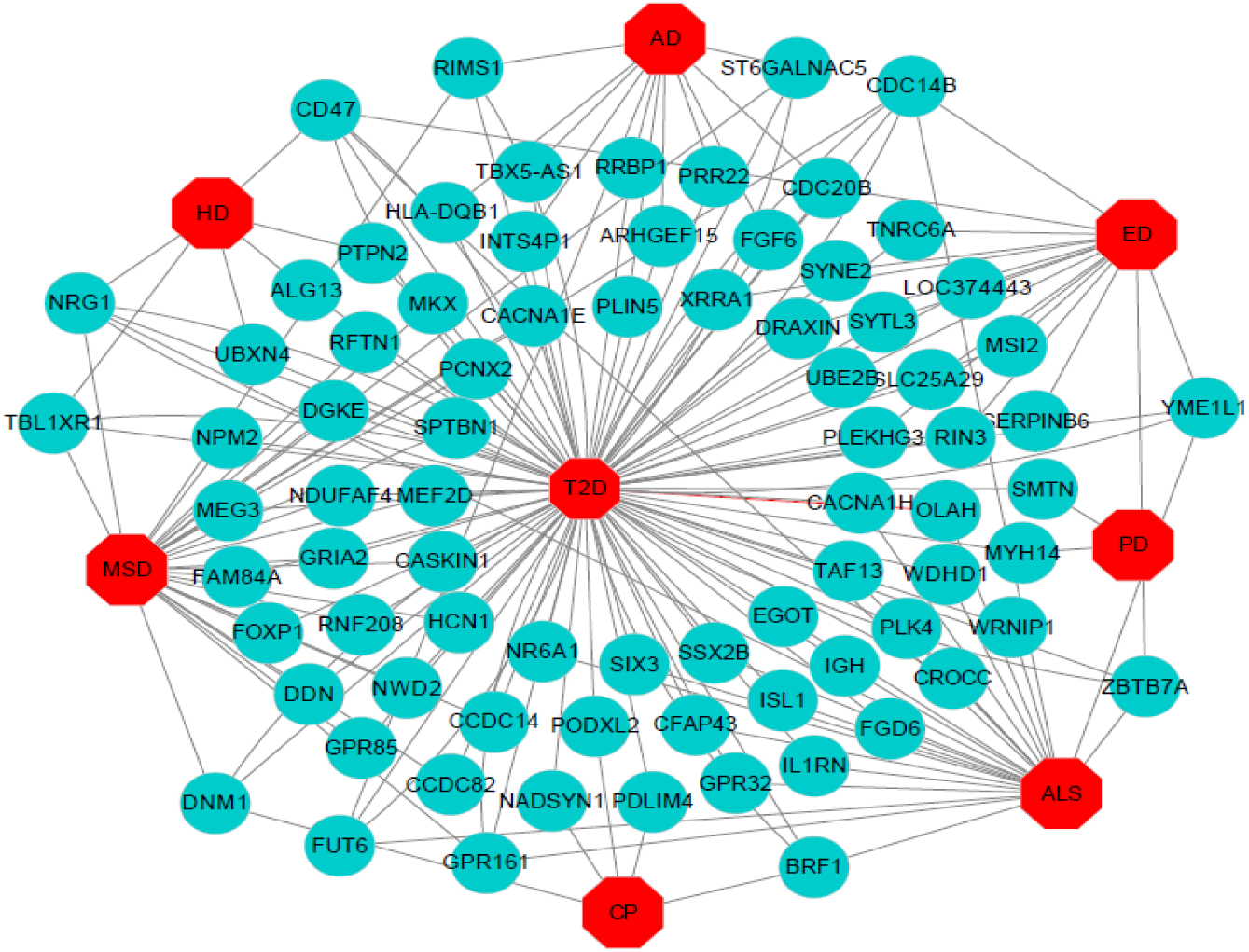
Disease network of T2D with NDs. Red colored octagon-shaped nodes represent different categories of disease, and round-shaped green colored nodes represent commonly down-regulated genes for T2D with the NDs. A link is placed between a disease and a gene if mutations in that gene lead to the specific disease.

### B. Pathway and Functional Association Analysis

Pathways are the key to know how an organism reacts to perturbations in its internal changes. The pathway-based analysis is a modern technique to understand how different complex diseases are related to each other by underlying molecular or biological mechanisms [35]. We analyzed pathway of the common differentially expressed genes using Enrichr[32], and KEGG pathway database [33] by determining T2D vs. AD, T2D vs. PD, T2D vs. ALS, T2D vs. ED, T2D vs. HD, T2D vs. CP, and T2D vs. MSD enrichment to identify overrepresented pathway groups amongst differentially expressed genes and to group them into functional categories.. Pathways deemed significantly enriched in the common DEG sets (FDR ¡i0.05) were reduced by manual curation to include only those with known relevance to the diseases concerned. The total numbers of 51 pathways data are summarized on Table I. We observe that 12,6,2,2,12,7, and 10 significant pathway, associated pathway genes, and adjusted P-values are identified by Enrichr which are significantly connected with DEG of T2D(See Table I: A, B, C, D, E, F, and G).

Moreover, We observed a number of the significant pathways that notably includes Synaptic vesicle cycle (hsa04721) among AD, CP, and MSD, Glycosphingolipid biosynthesis-ganglio series (hsa00604) between AD and MSD, Allograft rejection (hsa05330), Graft-versus-host disease (hsa05332), and Type I diabetes mellitus (hsa04940) between AD and PD, Glycosphingolipid biosynthesis-lacto and neolacto series (hsa00601) among ALS HD and AD, Purine metabolism (hsa00230) between CP and AD, Endocrine and other factor-regulated calcium reabsorption (hsa04961) between CP and MSD, Viral myocarditis (hsa05416) between HD and AD, and Arrhythmogenic right ventricular cardiomyopathy (ARVC) (hsa05412),Hypertronic cardiomyopathy(HCM) (hsa05410) and Dilated cardiomyopathy (hsa05414) between HD and MSD were observed.

**Table I:**
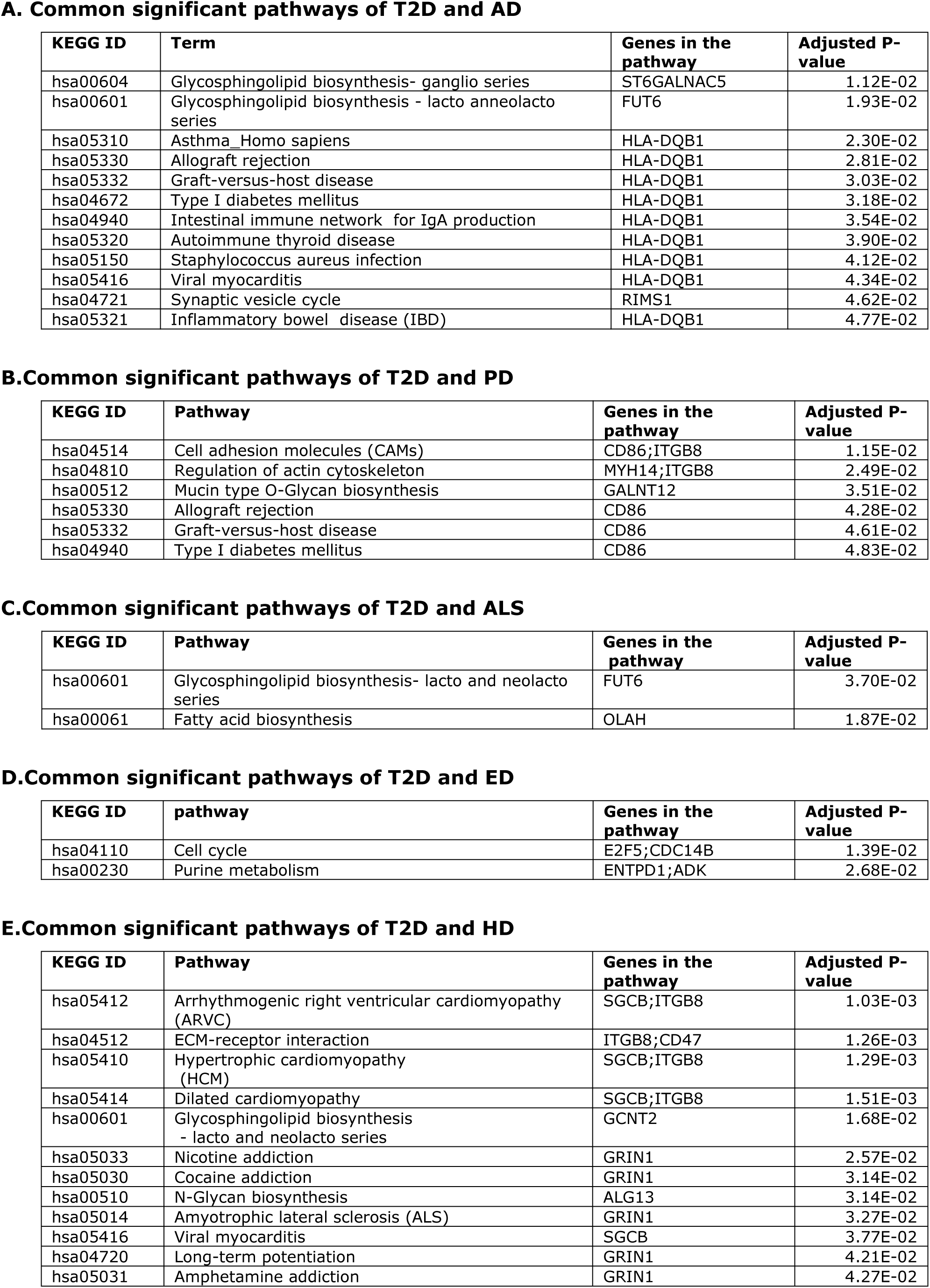
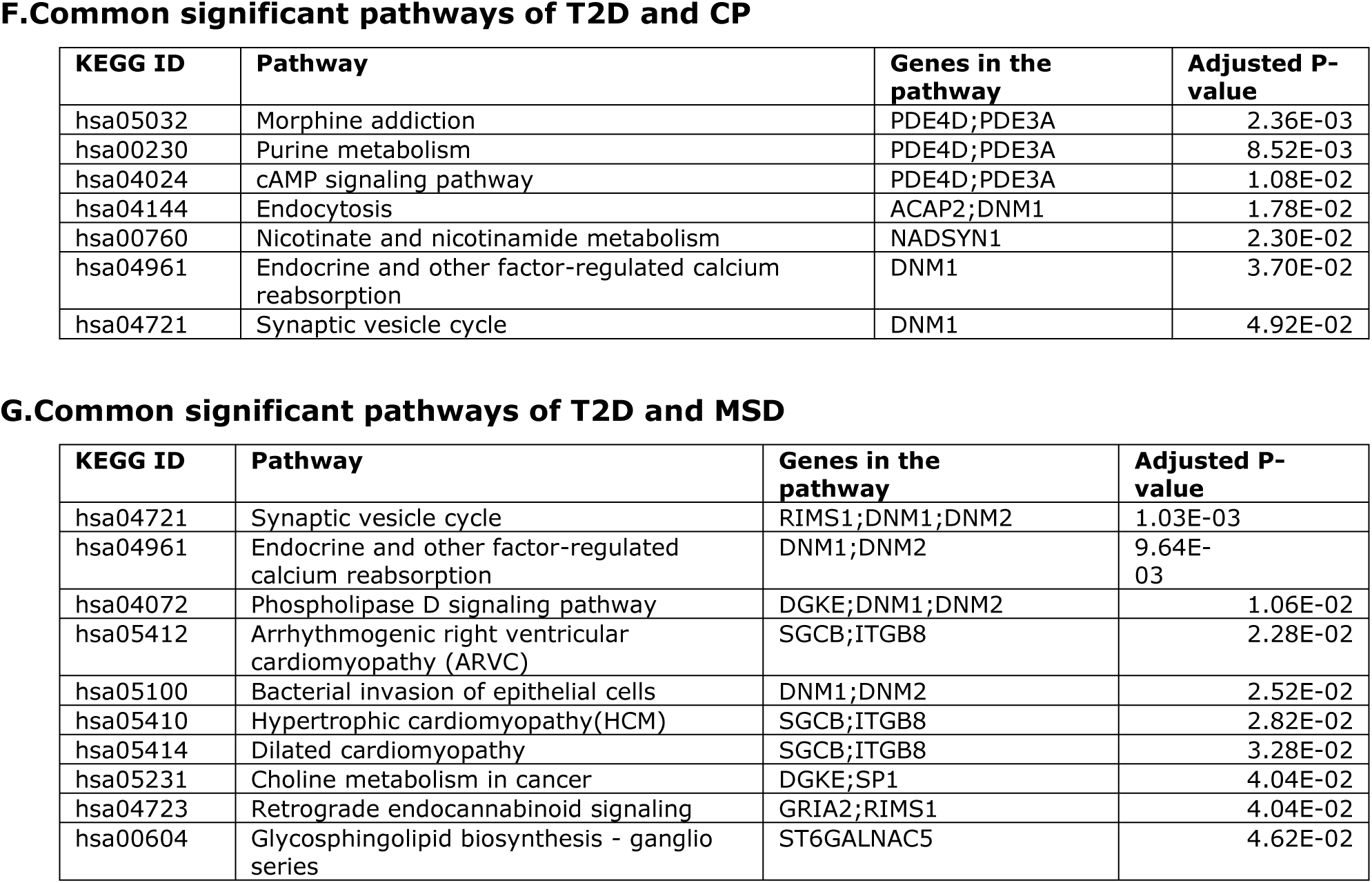
KEGG pathway analyses to identify pathways common to T2D and the NDs revealed by the commonly expressed genes. These include significant pathways common to T2D and A) AD B) PD C) ALS D) ED E) HD F) CP and G) MSD

A. Common significant pathways of T2D and AD
| KEGG ID | Term | Genes in the pathway | Adjusted P-value |
| --- | --- | --- | --- |
| hsa00604 | Glycosphingolipid biosynthesis- ganglio series | ST6GALNAC5 | 1.12E-02 |
| hsa00601 | Glycosphingolipid biosynthesis - lacto and neolacto series | FUT6 | 1.93E-02 |
| hsa05310 | Asthma_Homo sapiens | HLA-DQB1 | 2.30E-02 |
| hsa05330 | Allograft rejection | HLA-DQB1 | 2.81E-02 |
| hsa05332 | Graft-versus-host disease | HLA-DQB1 | 3.03E-02 |
| hsa04672 | Type I diabetes mellitus | HLA-DQB1 | 3.18E-02 |
| hsa04940 | Intestinal immune network for IgA production | HLA-DQB1 | 3.54E-02 |
| hsa05320 | Autoimmune thyroid disease | HLA-DQB1 | 3.90E-02 |
| hsa05150 | Staphylococcus aureus infection | HLA-DQB1 | 4.12E-02 |
| hsa05416 | Viral myocarditis | HLA-DQB1 | 4.34E-02 |
| hsa04721 | Synaptic vesicle cycle | RIMS1 | 4.62E-02 |
| hsa05321 | Inflammatory bowel disease (IBD) | HLA-DQB1 | 4.77E-02 |

B. Common significant pathways of T2D and PD
| KEGG ID | Pathway | Genes in the pathway | Adjusted P-value |
| --- | --- | --- | --- |
| hsa04514 | Cell adhesion molecules (CAMs) | CD86;ITGB8 | 1.15E-02 |
| hsa04810 | Regulation of actin cytoskeleton | MYH14;ITGB8 | 2.49E-02 |
| hsa00512 | Mucin type O-Glycan biosynthesis | GALNT12 | 3.51E-02 |
| hsa05330 | Allograft rejection | CD86 | 4.28E-02 |
| hsa05332 | Graft-versus-host disease | CD86 | 4.61E-02 |
| hsa04940 | Type I diabetes mellitus | CD86 | 4.83E-02 |

C. Common significant pathways of T2D and ALS
| KEGG ID | Pathway | Genes in the pathway | Adjusted P-value |
| --- | --- | --- | --- |
| hsa00601 | Glycosphingolipid biosynthesis- lacto and neolacto series | FUT6 | 3.70E-02 |
| hsa00061 | Fatty acid biosynthesis | OLAH | 1.87E-02 |

D. Common significant pathways of T2D and ED
| KEGG ID | pathway | Genes in the pathway | Adjusted P-value |
| --- | --- | --- | --- |
| hsa04110 | Cell cycle | E2F5;CDC14B | 1.39E-02 |
| hsa00230 | Purine metabolism | ENTPD1;ADK | 2.68E-02 |

E. Common significant pathways of T2D and HD
| KEGG ID | Pathway | Genes in the pathway | Adjusted P-value |
| --- | --- | --- | --- |
| hsa05412 | Arrhythmogenic right ventricular cardiomyopathy (ARVC) | SGCB;ITGB8 | 1.03E-03 |
| hsa04512 | ECM-receptor interaction | ITGB8;CD47 | 1.26E-03 |
| hsa05410 | Hypertrophic cardiomyopathy (HCM) | SGCB;ITGB8 | 1.29E-03 |
| hsa05414 | Dilated cardiomyopathy | SGCB;ITGB8 | 1.51E-03 |
| hsa00601 | Glycosphingolipid biosynthesis - lacto and neolacto series | GCNT2 | 1.68E-02 |
| hsa05033 | Nicotine addiction | GRIN1 | 2.57E-02 |
| hsa05030 | Cocaine addiction | GRIN1 | 3.14E-02 |
| hsa00510 | N-Glycan biosynthesis | ALG13 | 3.14E-02 |
| hsa05014 | Amyotrophic lateral sclerosis (ALS) | GRIN1 | 3.27E-02 |
| hsa05416 | Viral myocarditis | SGCB | 3.77E-02 |
| hsa04720 | Long-term potentiation | GRIN1 | 4.21E-02 |
| hsa05031 | Amphetamine addiction | GRIN1 | 4.27E-02 |

F.Common significant pathways of T2D and CP
| KEGG ID | Pathway | Genes in the pathway | Adjusted P-value |
| --- | --- | --- | --- |
| hsa05032 | Morphine addiction | PDE4D;PDE3A | 2.36E-03 |
| hsa00230 | Purine metabolism | PDE4D;PDE3A | 8.52E-03 |
| hsa04024 | cAMP signaling pathway | PDE4D;PDE3A | 1.08E-02 |
| hsa04144 | Endocytosis | ACAP2;DNM1 | 1.78E-02 |
| hsa00760 | Nicotinate and nicotinamide metabolism | NADSYN1 | 2.30E-02 |
| hsa04961 | Endocrine and other factor-regulated calcium reabsorption | DNM1 | 3.70E-02 |
| hsa04721 | Synaptic vesicle cycle | DNM1 | 4.92E-02 |

G.Common significant pathways of T2D and MSD
| KEGG ID | Pathway | Genes in the pathway | Adjusted P-value |
| --- | --- | --- | --- |
| hsa04721 | Synaptic vesicle cycle | RIMS1;DNM1;DNM2 | 1.03E-03 |
| hsa04961 | Endocrine and other factor-regulated calcium reabsorption | DNM1;DNM2 | 9.64E-03 |
| hsa04072 | Phospholipase D signaling pathway | DGKE;DNM1;DNM2 | 1.06E-02 |
| hsa05412 | Arrhythmogenic right ventricular cardiomyopathy (ARVC) | SGCB;ITGB8 | 2.28E-02 |
| hsa05100 | Bacterial invasion of epithelial cells | DNM1;DNM2 | 2.52E-02 |
| hsa05410 | Hypertrophic cardiomyopathy(HCM) | SGCB;ITGB8 | 2.82E-02 |
| hsa05414 | Dilated cardiomyopathy | SGCB;ITGB8 | 3.28E-02 |
| hsa05231 | Choline metabolism in cancer | DGKE;SP1 | 4.04E-02 |
| hsa04723 | Retrograde endocannabinoid signaling | GRIA2;RIMS1 | 4.04E-02 |
| hsa00604 | Glycosphingolipid biosynthesis - ganglio series | ST6GALNAC5 | 4.62E-02 |

### C. Gene Ontology Analysis

The Gene Ontology (GO) refers to a universal conceptual model for representing gene functions and their relationship in the domain of gene regulation. It is constantly expanded by accumulating the biological knowledge to cover regulation of gene functions and the relationship of these functions in terms of ontology classes and semantic relations between classes [36]. To get further insight into the identified pathways, enriched common gene sets were processed by gene ontology methods using Enrichr which identifies related biological processes and groups them into functional categories. The list of processes was also curated for those with known involvement with the diseases of interest. The cell processes and genes identified are summarized on Table II. We observe that 10,20,29,18,14,24, and 30 significant ontological pathways, associated pathway genes and adjusted P-values are identified by Enrichr which are significantly connected with DEGs of T2D(See table A, B, C, D, E, F, and G). Moreover,we observed a number of the significant pathways that notably includes regulation of neurotransmitter transport (GO:0051588) and regulation of neurotransmitter secretion (GO:0046928) between AD and MSD, negative regulation of nervous system development (GO:0051961) between ALS and AD, negative regulation of signal transduction (GO:0009968) between ALS and CP, central nervous system neuron differentiation (GO:0021953) between ALS and ED, positive regulation of cellular process (GO:0048522), neural fate commitment (GO:0048663) and neural fate specification (GO:0048665) between ALS and HD, regulation of ERK1 and ERK2 cascade (GO:0070372) among ALS, HD, and CP, neural crest cell differentiation (GO:0014033) among ALS, HD, and MSD, regulation of cell motility (GO:2000145) between ALS and MSD, neuronal action potential (GO:0019228) between ALS and PD, neurotransmitter receptor internalization (GO:0099590) between CP and MSD, protein localization to organelle (GO:0033365) between ED and MSD, mitotic spindle elongation (GO:0000022) between ED and PD, transmembrane receptor protein tyrosine kinase signaling pathway (GO:0007169) between HD and CP and finally extracellular matrix organization (GO:0030198) between HD and PD were observed.

**Table II:** Gene ontology identification of biological processes common to T2D and NDs.

A. Common significant GOs of T2D and AD
| GO ID | Pathway | Genes in the pathway | Adjusted P-value |
| --- | --- | --- | --- |
| GO:0032000 | positive regulation of fatty acid beta-oxidation | PLIN5 | 5.99E-03 |
| GO:0016064 | immunoglobulin mediated immune response | HLA-DQB1 | 6.73E-03 |
| GO:2000269 | regulation of fibroblast apoptotic process | STK17B | 6.73E-03 |
| GO:0031998 | regulation of fatty acid beta-oxidation | PLIN5 | 8.22E-03 |
| GO:0051588 | regulation of neurotransmitter transport | RIMS1 | 1.05E-02 |
| GO:0007498 | mesoderm development | GDF11 | 1.64E-02 |
| GO:0051961 | negative regulation of nervous system development | ARHGEF15 | 1.71E-02 |
| GO:0046928 | regulation of neurotransmitter secretion | RIMS1 | 2.08E-02 |
| GO:0043603 | cellular amide metabolic process | FUT6 | 4.62E-02 |
| GO:0009311 | oligosaccharide metabolic process | ST6GALNAC5 | 4.70E-02 |

B. Common significant GOs of T2D and PD
| GO ID | Pathway | Genes in the pathway | P-value |
| --- | --- | --- | --- |
| GO:0070486 | leukocyte aggregation | S100A8 | 9.16E-03 |
| GO:0070584 | mitochondrion morphogenesis | MYH14 | 1.14E-02 |
| GO:0090128 | regulation of synapse maturation | ADGRL1 | 1.26E-02 |
| GO:0045624 | positive regulation of T-helper cell differentiation | CD86 | 1.37E-02 |
| GO:0000022 | mitotic spindle elongation | CDC14A | 1.37E-02 |
| GO:1900003 | regulation of serine-type endopeptidase activity | CR1 | 1.37E-02 |
| GO:0032990 | cell part morphogenesis | MYH14 | 1.37E-02 |
| GO:0048854 | brain morphogenesis | SHANK2 | 1.37E-02 |
| GO:0006939 | smooth muscle contraction | SMTN | 1.71E-02 |
| GO:0003009 | skeletal muscle contraction | MYH14 | 2.16E-02 |
| GO:0043401 | steroid hormone mediated signaling pathway | NR6A1 | 2.39E-02 |
| GO:0019228 | neuronal action potential | MYH14 | 2.73E-02 |
| GO:0097061 | dendritic spine organization | SHANK2 | 2.84E-02 |
| GO:0030198 | extracellular matrix organization | DPT;ITGB8 | 2.84E-02 |
| GO:0030199 | collagen fibril organization | DPT | 3.40E-02 |
| GO:0061448 | connective tissue development | ITGB8 | 3.51E-02 |
| GO:0048813 | dendrite morphogenesis | SHANK2 | 3.73E-02 |
| GO:0009312 | oligosaccharide biosynthetic process | GALNT12 | 3.84E-02 |
| GO:0099601 | regulation of neurotransmitter receptor activity | SHANK2 | 3.84E-02 |
| GO:1901606 | alpha-amino acid catabolic process | MCCC2 | 4.50E-02 |

C.Common significant GOs of T2D and ALS
| GO ID | Pathway | Genes in the pathway | Adjusted P-value |
| --- | --- | --- | --- |
| GO:0043496 | regulation of protein homodimerization activity | NRG1;ISL1 | 2.72E-04 |
| GO:0072091 | regulation of stem cell proliferation | SIX3;FBLN1 | 2.72E-04 |
| GO:0045892 | negative regulation of transcription, DNA-templated | NR6A1;SIX3;NRG1;NR2F2;ZBTB7A;ISL1 | 9.49E-04 |
| GO:1902117 | positive regulation of organelle assembly | PLK4;CROCC | 2.20E-03 |
| GO:0006352 | DNA-templated transcription, initiation | NR6A1;BRF1;TAF13 | 2.46E-03 |
| GO:0050870 | positive regulation of T cell activation | TFRC;CD47 | 4.48E-03 |
| GO:2000113 | negative regulation of cellular macromolecule biosynthetic process | SIX3; NRG1; NR2F2; ZBTB7A | 6.11E-03 |
| GO:0048935 | peripheral nervous system neuron development | ISL1 | 1.01E-02 |
| GO:0021522 | spinal cord motor neuron differentiation | ISL1 | 1.01E-02 |
| GO:0021559 | trigeminal nerve development | ISL1 | 1.01E-02 |
| GO:2000145 | regulation of cell motility | NRG1;FBLN1 | 1.10E-02 |
| GO:0048665 | neuron fate specification | ISL1 | 1.15E-02 |
| GO:0051094 | positive regulation of developmental process | TFRC;ISL1 | 1.18E-02 |
| GO:1902692 | regulation of neuroblast proliferation | SIX3 | 1.58E-02 |
| GO:0048663 | neuron fate commitment | ISL1 | 2.01E-02 |
| GO:2000177 | regulation of neural precursor cell proliferation | SIX3 | 2.15E-02 |
| GO:0014033 | neural crest cell differentiation | NRG1 | 2.30E-02 |
| GO:0008284 | positive regulation of cell proliferation | NRG1;CD47;CYR61 | 2.31E-02 |
| GO:0006355 | regulation of transcription, DNA-templated | NR6A1;TAF13;SIX3;NRG1;NR2F2;ZBTB7A | 2.49E-02 |
| GO:1901990 | regulation of mitotic cell cycle phase transition | PLK4;CDC14B | 2.93E-02 |
| GO:0045597 | positive regulation of cell differentiation | ISL1;CYR61 | 3.23E-02 |
| GO:0051961 | negative regulation of nervous system development | GPR161 | 3.28E-02 |
| GO:0019228 | neuronal action potential | CACNA1H | 3.42E-02 |
| GO:0048522 | positive regulation of cellular process | NRG1;CD47;ISL1 | 3.87E-02 |
| GO:0021953 | central nervous system neuron differentiation | ISL1 | 4.68E-02 |
| GO:0000122 | negative regulation of transcription from RNA polymerase II promoter | NR6A1; NR2F2; ISL1 | 4.78E-02 |
| GO:0120031 | plasma membrane bounded cell projection assembly | FGD6;CDC14B | 4.78E-02 |
| GO:0050768 | negative regulation of neurogenesis | SIX3 | 4.82E-02 |
| GO:0070372 | regulation of ERK1 and ERK2 cascade | FBLN1; CYR61 | 4.99E-02 |

D.Common significant GOs of T2D and ED
| GO ID | Pathway | Genes in the pathway | Adjusted P-value |
| --- | --- | --- | --- |
| GO:0006897 | endocytosis | PACSIN2;RIN3;IGHA1 | 6.45E-03 |
| GO:0033365 | protein localization to organelle | PACSIN2;BICD1 | 8.85E-03 |
| GO:0051251 | positive regulation of lymphocyte activation | CD47;IGHA1 | 1.01E-02 |
| GO:0007599 | hemostasis | FLI1 | 1.15E-02 |
| GO:0015822 | ornithine transport | SLC25A29 | 1.15E-02 |
| GO:0090286 | cytoskeletal anchoring at nuclear membrane | SYNE2 | 1.30E-02 |
| GO:0034214 | protein hexamerization | YME1L1 | 1.44E-02 |
| GO:0007010 | cytoskeleton organization | PACSIN2;SYNE2 | 1.45E-02 |
| GO:0034063 | stress granule assembly | BICD1 | 1.73E-02 |
| GO:0000022 | mitotic spindle elongation | CDC14B | 1.73E-02 |
| GO:0097205 | renal filtration | IGHA1 | 1.73E-02 |
| GO:0035278 | miRNA mediated inhibition of translation | TNRC6A | 1.73E-02 |
| GO:0071470 | cellular response to osmotic stress | SERPINB6 | 2.15E-02 |
| GO:0034656 | nucleobase-containing small molecule catabolic process | ENTPD1 | 2.30E-02 |
| GO:0009069 | serine family amino acid metabolic process | SRR | 2.58E-02 |
| GO:0043949 | regulation of cAMP-mediated signaling | UBE2B | 2.72E-02 |
| GO:0072522 | purine-containing compound biosynthetic process | ADK | 3.28E-02 |
| GO:0021953 | central nervous system neuron differentiation | DRAXIN | 4.68E-02 |

E.Common significant GOs of T2D and HD
| GO ID | Pathway | Genes in the pathway | Adjusted P-value |
| --- | --- | --- | --- |
| GO:0042127 | regulation of cell proliferation | GCNT2;NRG1;CD47;POU3F2;PTPN2 | 6.92E-05 |
| GO:0022409 | positive regulation of cell-cell adhesion | GCNT2;CD47 | 2.16E-04 |
| GO:0048522 | positive regulation of cellular process | GCNT2;NRG1;CD47;POU3F2 | 2.68E-04 |
| GO:1901185 | negative regulation of ERBB signaling pathway | NRG1; PTPN2 | 3.63E-04 |
| GO:0051896 | regulation of protein kinase B signaling | GCNT2; NRG1 | 4.91E-03 |
| GO:0048665 | neuron fate specification | POU3F2 | 5.19E-03 |
| GO:0048663 | neuron fate commitment | POU3F2 | 9.06E-03 |
| GO:0030198 | extracellular matrix organization | ITGB8;CD47 | 9.45E-03 |
| GO:0014033 | neural crest cell differentiation | NRG1 | 1.04E-02 |
| GO:0070372 | regulation of ERK1 and ERK2 cascade | GCNT2;PTPN2 | 1.09E-02 |
| GO:0009968 | negative regulation of signal transduction | NRG1;PTPN2 | 1.41E-02 |
| GO:0043161 | proteasome-mediated ubiquitin-dependent protein catabolic process | TBL1XR1;UBXN4 | 1.49E-02 |
| GO:0007169 | transmembrane receptor protein tyrosine kinase signaling pathway | NRG1; PTPN2 | 2.65E-02 |
| GO:0007399 | nervous system development | NRG1;POU3F2 | 3.43E-02 |

F.Common significant GOs of T2D and CP
| GO ID | Pathway | Genes in the pathway | Adjusted P-value |
| --- | --- | --- | --- |
| GO:1903050 | regulation of proteolysis involved in cellular protein catabolic process | EPHA4;UBE3A | 1.64E-04 |
| GO:0046058 | cAMP metabolic process | PDE4D;PDE3A | 1.78E-04 |
| GO:0019933 | cAMP-mediated signaling | PDE4D;PDE3A | 1.65E-03 |
| GO:0048013 | ephrin receptor signaling pathway | EPHA4;DNM1 | 1.97E-03 |
| GO:0071363 | cellular response to growth factor stimulus | ACAP2;PDE3A | 5.47E-03 |
| GO:0048681 | negative regulation of axon regeneration | EPHA4 | 6.38E-03 |
| GO:0070571 | negative regulation of neuron projection regeneration | EPHA4 | 7.18E-03 |
| GO:0099590 | neurotransmitter receptor internalization | DNM1 | 8.77E-03 |
| GO:0048679 | regulation of axon regeneration | EPHA4 | 8.77E-03 |
| GO:0021952 | central nervous system projection neuron axonogenesis | EPHA4 | 8.77E-03 |
| GO:0008045 | motor neuron axon guidance | EPHA4 | 8.77E-03 |
| GO:0106030 | neuron projection fasciculation | EPHA4 | 8.77E-03 |
| GO:0009966 | regulation of signal transduction | EPHA4;IGFBP5 | 1.47E-02 |
| GO:0070372 | regulation of ERK1 and ERK2 cascade | EPHA4; P2RY1 | 1.64E-02 |
| GO:0071880 | adenylate cyclase-activating adrenergic receptor signaling pathway | PDE4D | 1.67E-02 |
| GO:0071875 | adrenergic receptor signaling pathway | PDE4D | 1.67E-02 |
| GO:1990090 | cellular response to nerve growth factor stimulus | ACAP2 | 1.82E-02 |
| GO:1901701 | cellular response to oxygen-containing compound | IGFBP5;PDE3A | 1.99E-02 |
| GO:0009968 | negative regulation of signal transduction | EPHA4;IGFBP5 | 2.11E-02 |
| GO:0043410 | positive regulation of MAPK cascade | EPHA4; P2RY1 | 2.20E-02 |
| GO:0007169 | transmembrane receptor protein tyrosine kinase signaling pathway | EPHA4; DNM1 | 3.93E-02 |
| GO:0010977 | negative regulation of neuron projection development | EPHA4 | 4.08E-02 |
| GO:0016192 | vesicle-mediated transport | ACAP2;DNM1 | 4.18E-02 |
| GO:0001934 | positive regulation of protein phosphorylation | EPHA4;P2RY1 | 4.22E-02 |

G.Common significant GOs of T2D and CP
| GO ID | Pathway | Genes in the pathway | Adjusted P-value |
| --- | --- | --- | --- |
| GO:0070229 | negative regulation of lymphocyte apoptotic process | PRKCQ; FOXP1 | 4.32E-04 |
| GO:0099590 | neurotransmitter receptor internalization | DNM1; DNM2 | 5.27E-04 |
| GO:0061025 | membrane fusion | RIMS1;DNM1;DNM2 | 6.95E-04 |
| GO:0043112 | receptor metabolic process | ACKR3;DNM1;DNM2 | 8.54E-04 |
| GO:0051588 | regulation of neurotransmitter transport | RIMS1; KCNMB4 | 8.67E-04 |
| GO:0050804 | modulation of chemical synaptic transmission | GRIA2;RIMS1;KCNMB4 | 2.29E-03 |
| GO:0046928 | regulation of neurotransmitter secretion | RIMS1;KCNMB4 | 3.50E-03 |
| GO:2000146 | negative regulation of cell motility | IGFBP5;FBLN1;SRGAP1 | 3.66E-03 |
| GO:0071320 | cellular response to cAMP | IGFBP5;HCN1 | 4.28E-03 |
| GO:0072384 | organelle transport along microtubule | TRAK1;BICD1 | 4.56E-03 |
| GO:0007009 | plasma membrane organization | PACSIN2;SPTBN1 | 6.06E-03 |
| GO:0048661 | positive regulation of smooth muscle cell proliferation | MEF2D;FOXP1 | 6.38E-03 |
| GO:0090181 | regulation of cholesterol metabolic process | SQLE;SP1 | 7.75E-03 |
| GO:0048660 | regulation of smooth muscle cell proliferation | IGFBP5;FOXP1 | 8.86E-03 |
| GO:0001508 | action potential | KCNMB4;CACNA1E | 1.30E-02 |
| GO:0007411 | axon guidance | FRS2;PRKCQ;SPTBN1 | 1.38E-02 |
| GO:0072657 | protein localization to membrane | PIGR;RFTN1;SPTBN1 | 1.43E-02 |
| GO:0007517 | muscle organ development | MKX;MEF2D | 1.59E-02 |
| GO:0007005 | mitochondrion organization | TRAK1;DNM1;DNM2 | 1.60E-02 |
| GO:0010595 | positive regulation of endothelial cell migration | SP1;FOXP1 | 2.11E-02 |
| GO:0010646 | regulation of cell communication | GRIA2;CXORF36 | 2.17E-02 |
| GO:0038127 | ERBB signaling pathway | PIGR;NRG1 | 2.70E-02 |
| GO:0050658 | RNA transport | RFTN1;SRSF3 | 2.70E-02 |
| GO:0007088 | regulation of mitotic nuclear division | NPM2;CDC14B | 2.95E-02 |
| GO:0010628 | positive regulation of gene expression | RIMS1;SP1;TBL1XR1<br>ITGB8;FLI1;DNM2 | 3.43E-02 |
| GO:0033365 | protein localization to organelle | PACSIN2;BICD1 | 3.83E-02 |
| GO:0098916 | anterograde trans-synaptic signaling | GRIA2;KCNMB4;CACNA1E | 4.05E-02 |
| GO:0006406 | mRNA export from nucleus | SRSF3;NUP58 | 4.41E-02 |
| GO:2000145 | regulation of cell motility | FBLN1;NRG1 | 4.71E-02 |
| GO:0014033 | neural crest cell differentiation | NRG1 | 4.92E-02 |

### D. Protein-protein-protein interaction (PPI) analysis

Protein-protein interaction networks (PPINs) are the mathematical representation of the physical contacts of proteins in the cell. Protein-protein interactions (PPIs) are essential to every molecular and biological process in a cell, so PPIs is crucial to understand cell physiology in disease and healthy states [37]. The malfunction of a protein complex causes multiple diseases by the malfunction of a protein complex. Two diseases are potentially related to each other if they share one or more commonly associated protein sub network. Having identified genes involved in pathways and processes common to T2D and the Neurological diseases, we sought evidence for existing sub-networks based on known PPI. Using the enriched common disease genesets, we constructed putative PPI networks using web-based visualization resource STRING [38] by the distinct 159 differentially expressed genes as shown in Fig3. The network is also grouped into 7 clusters using the MCL clustering technique representing NDs to depict the protein belongings. The CDC14B protein belongs to the maximum 3 clusters ALS, ED, and MSD whereas NRG1 protein belongs to the 3 clusters ALS, HD, and MSD which interacts with other proteins from different clusters.FLI1, PACSIN2, ENTPD1, ZBTB7A, DNM1, SGCB, and FUT6 proteins belong to two clusters and interact with proteins in the network. For topological analysis, a simplified PPI network was constructed using Cyto-Hubba plugin [39] to show 10 most significant hub proteins as shown in Fig. 4 which are DNM2, DNM1, MYH14, PACSIN2, TFRC, PDE4D, ENTPD1, PLK4, CDC20B, and CDC14A. This data provides evidence that PPI sub-network exists in our enriched genesets, and confirm the presence of relevant functional pathways. Moreover, thus, these proteins could be the targeted proteins for drug development.

**Fig 3.**
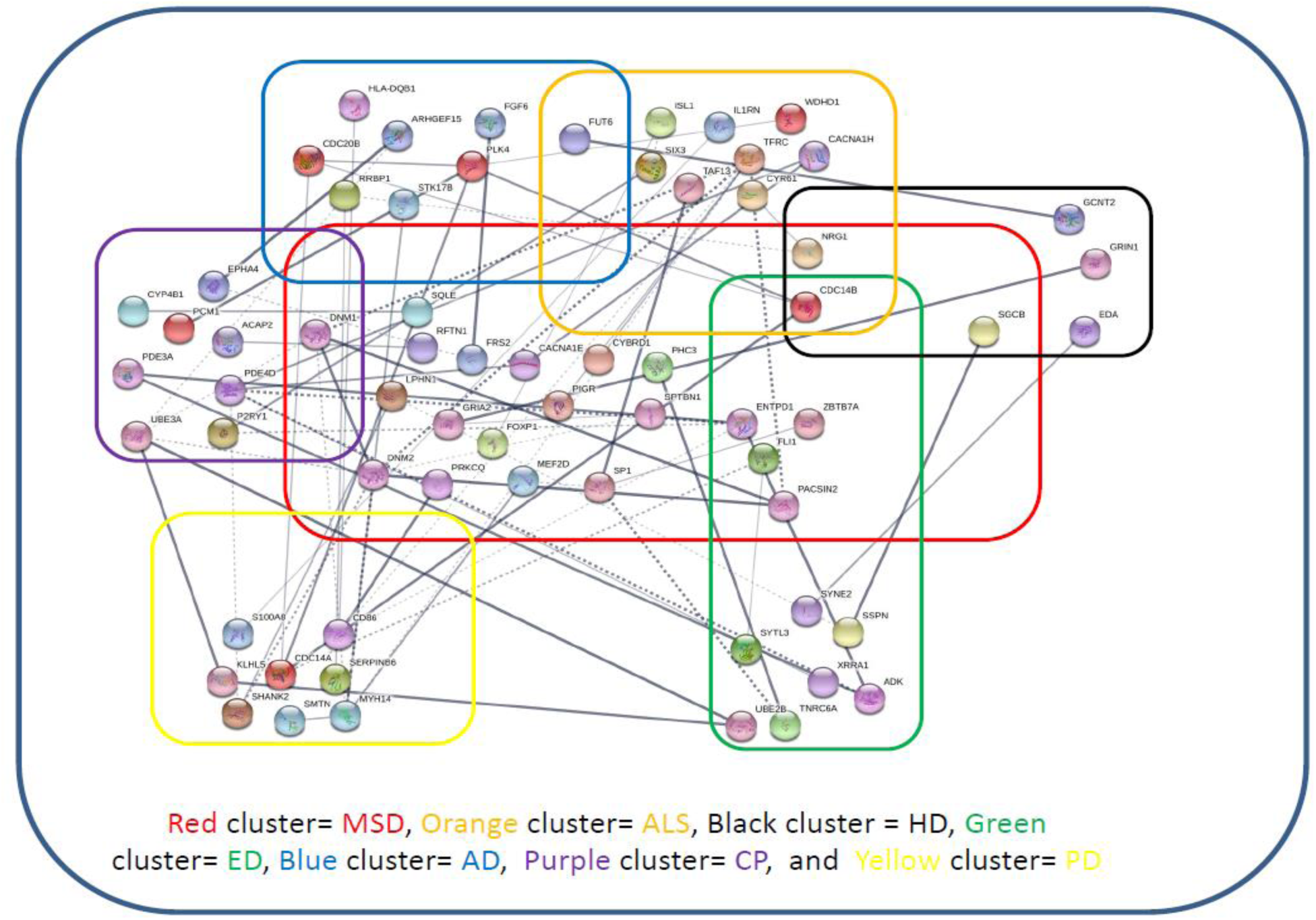
Protein-protein interaction network of the commonly significant Dysregulated genes of the NDs with T2D. Each cluster indicates the gene belongings.

**Fig 4.**
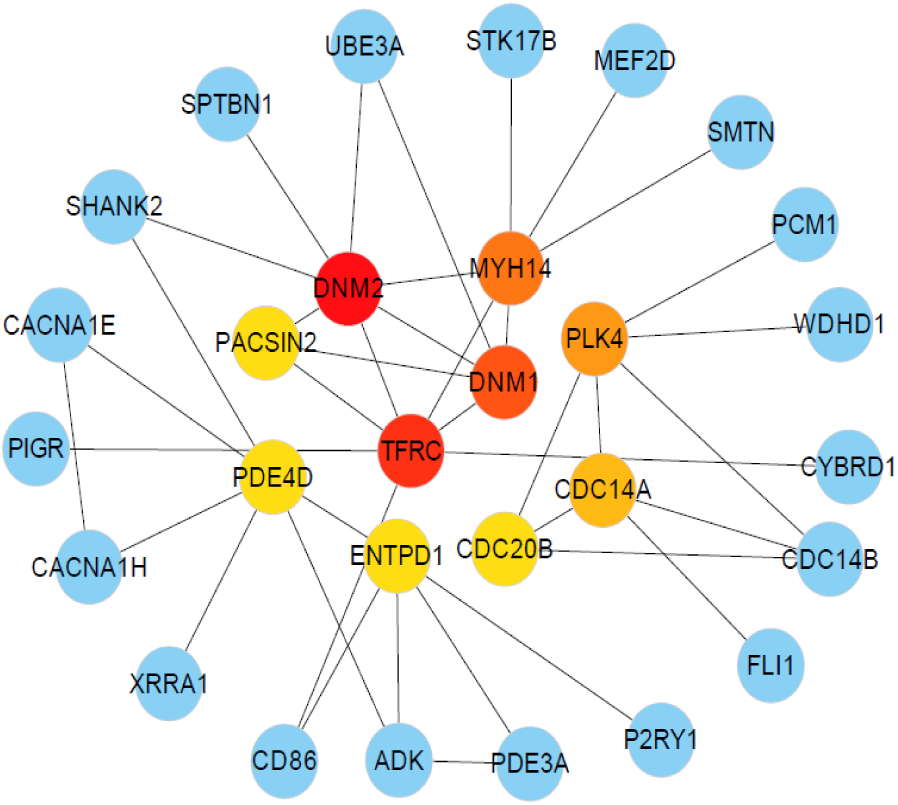
The simplified PPI network of the commonly dysregulated genes between NDs and T2D. The 10 most significant hub proteins are marked as red,orange and yellow.

### E. Validating Biomarkers by Gold Benchmark Databases

We presented a combined relation of OMIM, OMIM Expanded and dbGap databases. For cross checking the validity of our study, we collected genes and disease names from OMIM Disease, OMIM Expanded and dbGap databases using differentially expressed genes of T2D. To find out significant neurological diseases, manual curation is applied considering adjusted p-value below or equal to 0.05. Then, several diseases such as cancer, infectious diseases etc. are removed from this list because they are not concerned in this study. After analyzing them, 5 neurological diseases are found. Then, we construct a GDN using Cytoscape and show gene-disease association of different diseases as shown in Fig. 5. It indicates our analysis of finding significant genes of neurological diseases are also matched with existing records.

**Fig 5.**
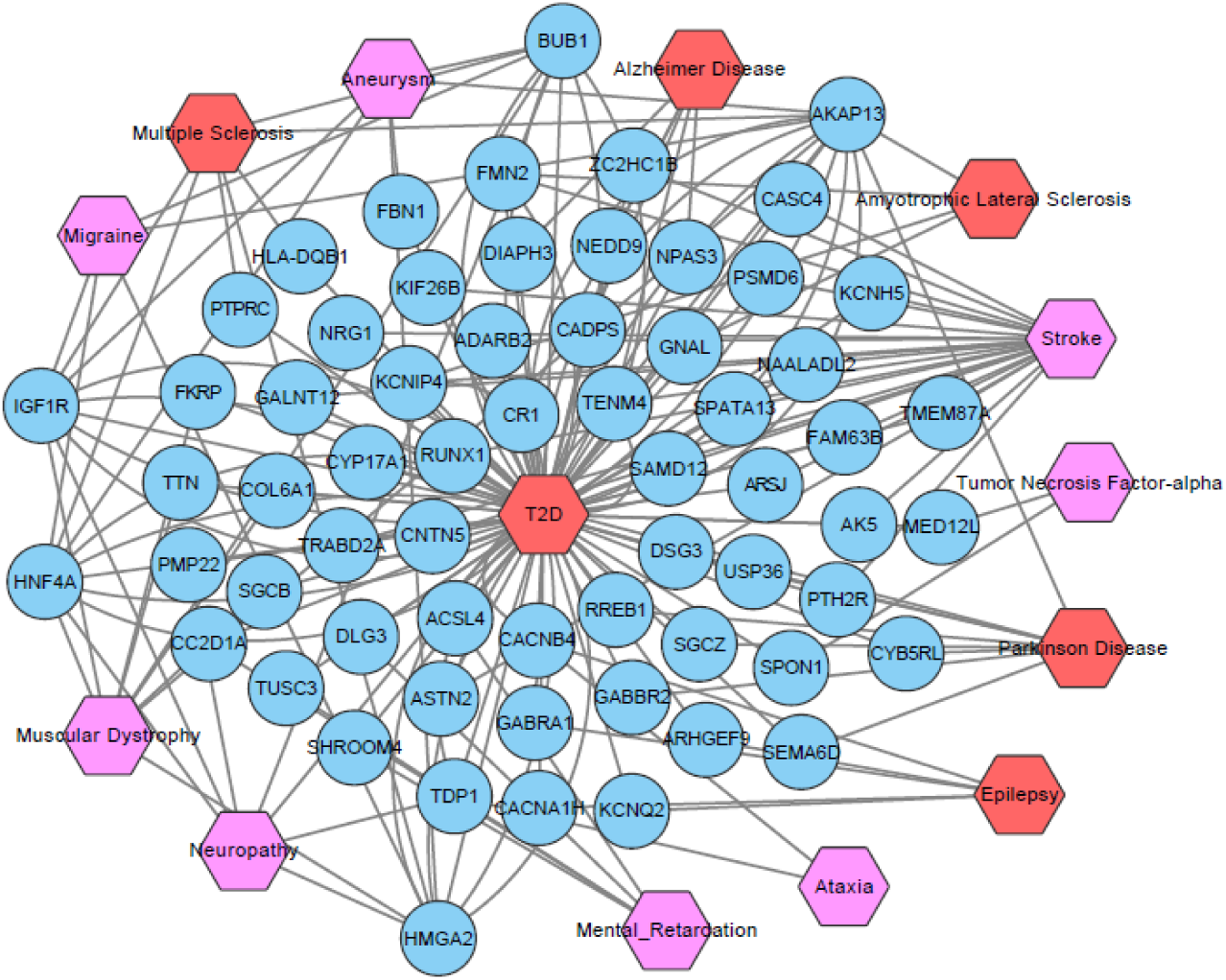
Disease network of T2D with several NDs.Red color hexagon-shaped nodes represent our selected five NDs. Violet colored hexagon-shaped nodes represent different categories of NDs. A link is placed between a disease and gene if muttions inthat gene lead to the specific diseases.

## IV. DISCUSSION

Our analysis fills a significant gap in our knowledge about how T2D may affect AD, PD, ALS, ED, HD, CP, and MSD. We used a simple sequence of steps that employ widely available resources and data and can easily be applied to a range of possible co-morbidities. Based on the combined analysis of transcriptomics, genetics, PPIs, pathways and GO data, our disease network disclosed potentially novel disease relationships that have not been captured by previous individual studies. These may inform future clinical co-morbidity studies. The underlying hypothesis behind this line of research is that once we catalogue all (or a large proportion of) disease-related genes, PPI complex, and signaling pathways, we will be able to predict the susceptibility of individuals to other diseases using molecular biomarkers. Combined with genetic data such analyses will be a key element in the development of truly predictive medicine. Our results show a combination of molecular and population-level data that provide insights about the novel hypothesis of disease mechanism. Furthermore, it will provide important information about medication overlaps and the probability of developing disease co-morbidity, i.e., where the occurrence of one disease in a patient may increase the susceptibility to or severity of another disease using molecular biomarkers. This observation makes us to understand the correlation between various neurological diseases from the molecular and genetic aspects. This analysis will also be considered as a key element of predictive drugs development.

This work showed significant associations of neurological diseases with the corresponding T2D by substantial pathways. To explore the pathway (and transcript profile) overlaps can indicate disease associations and co-morbid vulnerability. We analyzed publicly available microarray data thus can be applied wherever such data exists. Our approach employed differentiation gene expression analyses, followed by gene enrichment using signaling pathway and Gene Ontology (GO) data. One particular technical point is that with pathway and GO analyses a large number of categories were reduced by manual curation. Besides, flexible, time-consuming and semantic analysis-based approaches are used to facilitate this work and reduce operator bias. In transcript analyses, it is found evidence about the processing of disease of PPI data. So, we can identify different pathways through the inspection of cell proteins and their interactions. This investigation also represents a high potential for understanding the central mechanisms behind the disease or disorder progression. We have also analyzed the differentially expressed genes of T2D with gold benchmark dataset OMIM, OMIM Expanded and dbGaP databases using EnrichR to validate our identified results and depicted as gene-disease associations as shown in Fig. 4. These results corroborate that, the differentially expressed genes of T2D are responsible for the progression of NDs. As a whole, our findings compensate a major gap of about T2D biology. It will also open up an entry point to establish a mechanic link between the T2D and various neurological diseases.

## V. CONCLUSIONS

In this study, we have considered Gene Expression Omnibus (GEO) microarray data from type 2 diabetes (T2D), Parkinson’s disease (PD), Alzheimer’s disease (AD), Amyotrophic Lateral Sclerosis(Lou Gehrig’s disease) ALS (LGD), Epilepsy disease (ED),Huntington Disease (HD),Cerebral Palsy(CP), Multiple Sclerosis disease (MSD) and control datasets to analysis and investigate the genetic effects of T2D on neurological diseases (NDs). We analyzed dysregulated genes, disease relationship networks, dysregulated pathways, gene expression ontologies and protein-protein interactions of T2D and NDs. Our findings showed that T2D have a strong association with NDs. This study demonstrates that T2D shares several common multifactorial degenerative biological process that contributes to neuronal death,which leads to functional impairment. Because of these multifactorial aspects and complexity, our proposed gene expression analysis platforms have been extensively used to investigate altered pathways and to identify potential biomarkers and drug targets. Although many therapeutic approaches has been tested, no effective cure for these neurological diseases has been identified. Therefore, high-throuhput technique like whole genome transcriptomics and microarray technology must be coupled with functional genomics and proteomics in an effort to identify specific and selective biomarkers and viable drug targets which allow the successful discovery of disease modifying therapeutic targets. Therapeutic targets aimed at attenuating the above mentioned altered pathways could possibly ameliorate neurological dysfunction in T2D patient. This kind of study will be useful for making genomic evidence-based recommendations about the accurate disease prediction, identification, and making society aware of the dangerous effect of T2D on the human body.

## Acknowledgments

Thanks are due to Dou Hao and those anonymous scholars who offered advice on the improvement of the paper. This work is financially supported by the National Natural Science Foundation of China (Grant No.61571438). Thanks to CAS-TWAS Presidents Fellowship for supporting me(MHR) as a doctoral student (Fellowship No.2016CTF0146).

